# TET-mediated epimutagenesis of the *Arabidopsis thaliana* methylome

**DOI:** 10.1101/151027

**Authors:** Lexiang Ji, William T. Jordan, Xiuling Shi, Lulu Hu, Chuan He, Robert J. Schmitz

## Abstract

DNA methylation in the promoters of plant genes sometimes leads to transcriptional repression, and the wholesale removal of DNA methylation as seen in methyltransferase mutants results in drastic changes in gene expression and severe developmental defects. However, many cases of naturally-occurring DNA methylation variations have been reported, whereby the altered expression of differentially methylated genes is responsible for agronomically important traits. The ability to manipulate plant methylomes to generate populations of epigenetically distinct individuals could provide invaluable resources for breeding and research purposes. Here we describe “epimutagenesis”, a novel method to rapidly generate variation of DNA methylation through random demethylation of the *Arabidopsis thaliana* genome. This method involves the expression of a human Ten-eleven translocation (TET) enzyme, and results in widespread hypomethylation that can be inherited to subsequent generations, mimicking mutants in the maintenance DNA methyltransferase *met1*. Application of TET-mediated epimutagenesis to agriculturally significant plants may result in differential expression of alleles normally silenced by DNA methylation, uncovering previously hidden phenotypic variations.

---

Our ability to develop novel beneficial crop traits has significantly improved over the last 100 years, although the ability to maintain this trajectory is limited by allelic diversity. While genetic variation has been heavily exploited for crop improvement, utility of epigenetic variation has yet to be efficiently implemented. Epigenetic variation arises not from a change in the DNA sequence, but by changes in modifications to DNA such as cytosine methylation. This variation can result in emergence of novel and stably inherited phenotypes, as well as unique patterns of gene expression.

In plant genomes, cytosine methylation occurs at three major sequence contexts: CG, CHG and CHH (where H = A, C or T)^1^. Methylation at these different contexts is coordinated by distinct maintenance mechanisms during DNA replication. The methylation of DNA in all three contexts is essential for transcriptional silencing of transposons, repeat sequences and certain genes. Genes regulated by this mechanism are stably repressed throughout the soma and represent an untapped source of hidden genetic variation if transcriptionally re-activated, as revealed from pioneering studies in the model plant *A. thaliana*^2, 3, 4^. However, the impact of this variation is not observed in wild-type plants, as genes silenced by DNA methylation are not expressed. This novel source of genetic variation was uncovered by creating epigenetic recombinant inbred lines (epiRILs) from crosses between a wild-type individual and a mutant defective in maintenance of DNA methylation^2, 3, 4^. EpiRILs, while genetically wild type, contain mosaic DNA methylomes dependent on chromosomal inheritance patterns, as DNA methylation is meiotically inherited in *A. thaliana*^2, 5, 6, 7, 8^. Phenotypic characterization of epiRILs has revealed extensive morphological variation with respect to traits such as flowering time, root length and resistance to bacterial infection^2, 3, 4^. The morphological variation generated by the creation of epiRILs has revealed extensive hidden genetic variation in plant genomes that can be observed due to expression of newly unmethylated regions. However, the creation of epiRILs requires one founding parent to be a null mutant in the maintenance DNA methylation pathway. Unfortunately, unlike in *A. thaliana*, the loss of DNA methylation maintenance activity often results in lethality in crops^9, 10^. Therefore, novel methodologies are required to realize the potential of these hidden epialleles in crop genomes.

Epimutagenesis is an alternative method to generate epiRILs. Instead of relying on the genome-wide demethylation of one of the two founding parents, epimutagenesis introduces random methylation variation via the introduction of a transgene. Here, we describe a novel epimutagenesis approach in *A. thaliana* using a human Ten-eleven translocation (TET1) methylcytosine dioxygenase^11, 12, 13, 14^, which catalyzes the conversion of 5-methylcytosine (5mC) to 5-hydroxymethylcytosine (5hmC). Although TET enzymes and their primary product 5hmC are not found in plant genomes^15^, ectopic expression of a human TET enzyme resulted in widespread DNA demethylation and induced phenotypic variation in *A. thaliana*.

## RESULTS

### Overexpressing hTET1cd in Arabidopsis hypomethylates the genome

Transgenic *A. thaliana* plants were generated expressing the catalytic domain (residues 1418-2136) of the human TET1 protein (hTET1cd) under the control of the CaMV35S promoter. To assess the impact of hTET1cd expression on the *A. thaliana* methylome, whole genome bisulfite sequencing (WGBS) was performed on two independently derived transgenic plants (35S:TET1-1 and 35S:TET1-2; **Supplementary Table 1**). WGBS revealed a global reduction of CG methylation from 18.2% in two wild-type individuals to 8.9% in 35S:TET1-1 and 6.9% in 35S:TET1-2 (compared to 0.5% in *met1*-3). The effects of hTET1cd expression on CHG and CHH methylation were not as severe compared to CG methylation (Fig. 1a). Importantly, different degrees of CG hypomethylation were observed in different independent transgenic plants. This result has important implications for epimutagenesis in economically and agriculturally significant plant species, as it appears feasible to control the degree of DNA hypomethylation by screening for plants with desired levels of demethylation. Taken together, these results show that the expression of hTET1cd results in intermediate CG methylation levels when compared to wild-type and *met1* individuals.

**Figure 1.**
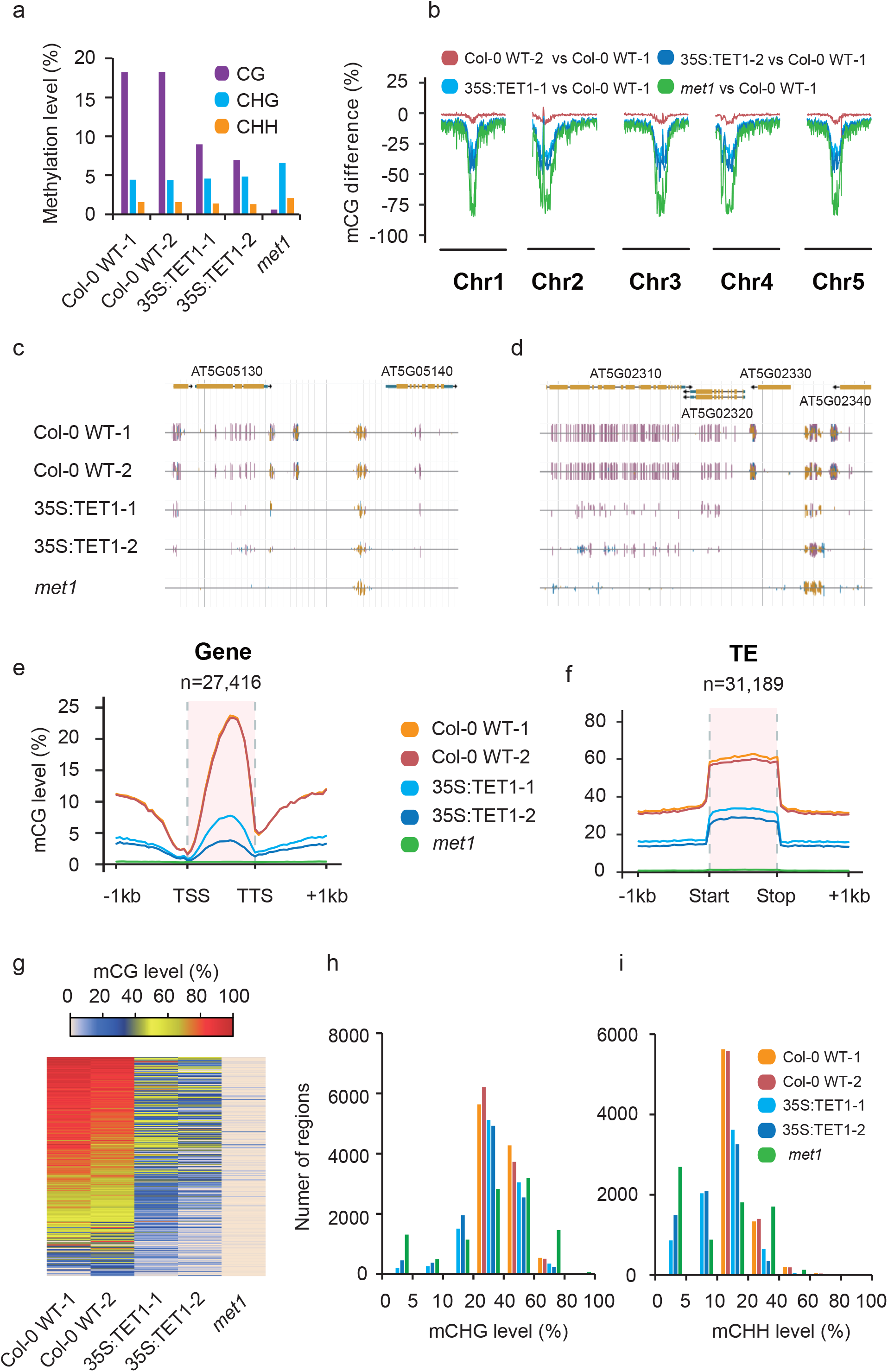
Overexpression of hTET1cd induced global CG demethylation in *A. thaliana*. (a) Bar graph of global methylation levels in two Col-0 WT replicates, two 35S:TET1 transgenic individuals and *met1*. (b) Metaplot of CG methylation levels (100 kb windows) across five *A. thaliana* chromosomes. Methylation level differences were defined relative to Col-0 WT-1, and Col-0 WT-2 was used to assess background interference. Genome browser view of methylome profile of two regions (c and d) of the *A. thaliana* genome (purple vertical lines = CG methylation, blue vertical lines = CHG methylation and gold vertical lines = CHH methylation). Metagene plots of CG methylation level across (e) gene bodies and (f) transposable elements. (g) Heat map of CG methylation level of CG DMRs. Bar plots of CHG (h) and CHH (i) methylation levels of CG DMRs that possess non-CG methylation in wild-type individuals.

The primary product of TET1 oxidation is 5hmC, which is indistinguishable from 5mC by WGBS. We therefore performed Tet-assisted bisulfite sequencing (TAB-seq) to profile 5hmC levels in 35S:TET1 plants^16^. No detectable levels of 5hmC were found in the transgenic lines assayed (**Supplementary Fig. 1a-b**). Thus, the widespread loss of CG DNA methylation observed may result from a failure to maintain methylation at CG sites that possess 5hmC, or through active removal of 5hmC or further oxidized products via the base excision repair pathway.

To better understand the effects of hTET1cd expression, we determined changes in the *A. thaliana* methylome at the chromosomal and local levels. Plotting methylation levels across all five chromosomes revealed a strong depletion of CG methylation at the pericentromeric region (Fig. 1b). CG hypomethylation occurred at both gene body methylated (gbM) and select RNA-directed DNA methylated (RdDM) loci. (Fig. 1c **and** d). To further quantify the observed hypomethylation, metaplots were created for genes and transposons, respectively (Fig. 1e **and** f, **Supplementary Fig. 1c-f**). Strong reduction of mCG and a mild reduction of mCHG/mCHH were observed at both genes and transposons. On average, 97.9% of gbM genes and 56.7% of methylated transposons (where these regions have at least 50% mCG in wild type) lost at least half of their CG methylation in epimutagenized lines (**Supplementary Table 5 and 6**). Collectively, these results indicate hypomethylation was more severe in genes than transposons, possibly the result of *de novo* methylation by the RdDM pathway which is primarily active at transposons.

### Tet1-mediated DNA demethylation mimics *met1* mutants

An analysis of Differentially Methylated Regions (DMRs) was then carried out to assess the genome-wide impact of hTET1cd expression. 56,283 CG DMRs ranging in size from 6 - 20,286 base pairs (bp) were identified (Fig. 1g, **Supplementary Table 7**). Of these, 38.7% were located in intergenic sequences, 53.7% overlapped with genes and 7.6% were located in promoter regions (≤ 1kb upstream of a gene). As also seen in *met1* mutants, the predominant effect of hTET1cd expression is CG hypomethylation (12,641 and 20,601 DMRs lost more than 50% mCG in 35S:TET1-1 and 35S:TET1-2, respectively; no region gained more than 50% mCG). However, the extent of CG methylation loss caused by hTETcd expression is lower than in *met1:* 31.8 Mb of the genome significantly lost CG methylation in *met1*, whereas 9.9 Mb and 18.0 Mb were lost in 35S:TET1-1 and 35S:TET1-2, respectively.

Previous studies of the *met1* methylome have revealed a loss of mCHG/mCHH methylation in a subset of CG-hypomethylated regions ^17^. At these loci, DNA methylation is stably lost, in contrast to regions where DNA methylation is re-established by *de novo* methylation pathways. These loci are ideal targets of epimutagenesis, as the co-existence of all three types of methylation is more frequently correlated with transcriptional repression of genes than CG methylation alone. This, coupled with the long-term stability of hypomethylation, may facilitate inherited transcriptional changes.

An analysis of the interdependence of the loss of CG methylation on non-CG methylation levels revealed that 39.7 Kb and 931.5 Kb of CHG methylated sequences lost significant amounts of methylation in two independent epimutagenized lines, compared to 4.0 Mb of sequence in *met1* mutants. A similar analysis for the loss of CHH methylation revealed losses of 23.3 Kb and 492.5 Kb in epimutagenized individuals, compared to 1.1 Mb lost in *met1* mutants. Of the 56,283 identified CG DMRs, 10,491 overlapped regions that contained at least 20% CHG methylation and 7,214 overlapped regions that contained at least 10% CHH methylation in wild-type individuals (**Supplementary Table 7**). To determine how many of these regions are susceptible to losing non-CG methylation if CG methylation is first depleted, we created a frequency distribution of mCHG and mCHH levels in wild-type and epimutagenized individuals (Fig. 1h and i). 2,341 and 3,447 regions lost more than 10% CHG methylation in 35S:TET1-1 and 35S:TET1-2, respectively, whereas 2,475 and 3,379 regions lost more than 5%

CHH methylation in 35S:TET1-1 and 35S:TET1-2, respectively. Regions that are susceptible to losses of CG and non-CG methylation in lines expressing hTET1cd share a substantial overlap with regions that lose non-CG methylation in *met1* (**Supplementary Fig. 1g and h**). 1,708 (73.0%) and 2,386 (69.2%) regions that have lost more than 10% mCHG in 35S:TET1-1 and 35S:TET1-2 have reduced levels in *met1*, whereas 2,013 (81.6%) and 2,563 (75.9%) regions that have lost more than 5% mCHH in 35S:TET1-1 and 35S:TET1-2 have reduced levels in *met1*. As crop genomes have a greater number of loci targeted for silencing by CG, CHG and CHH methylation when compared to *A. thaliana*, ectopic expression of hTET1cd is likely a viable approach for the creation epiRILs^18^.

### Tet1-mediated variation of CHG methylation

Mutations in *met1* also result in hypermethylation of CHG sites in gene bodies due to the loss of methylation in the 7^th^ intron of the histone 3 lysine 9 (H3K9) demethylase, *INCREASE IN BONSAI METHYLATION 1* (*IBM1*)^19, 20, 21, 22^. This results in alternative splicing of *IBM1*, ultimately producing a non-functional gene product (IBM1-S), which results in ectopic accumulation of di-methylation of H3K9 (H3K9me2) throughout the genome^23^. As in *met1*, the 7^th^ intron of *IBM1* was hypomethylated in 35S:TET1-1, 35S:TET1-2, and an additional two lines, 35S:TET1-2^T5^ and 35S:TET1-2^T6^, which were propagated for an additional two and three generations, respectively (Fig. 2a). The increased abundance of *IBM1-S* and the reduction of functional *IBM1 (IBM1-L)* transcript was confirmed by RT-qPCR in line 35S:TET1-2^T6^ (**Supplementary Fig. 2a**) leading to CHG hypermethylation at gbM loci (Fig. 2b). Further quantitative analysis revealed extensive variation in genome-wide gains and losses of CHG methylation in these two lines, approximately 1.8 Mb and 2.3 Mb of additional CHG methylation, respectively (Fig. 2c, **Supplementary Fig. 2b and c**). To test the impact of a reduction in functional *IBM1* on H3K9me2, we performed chromatin immunoprecipitation (ChIP) against H3K9me2 in 35S:TET1-2^T6^, which revealed a subtle increase in H3K9me2 in gbM loci that possessed CHG hypermethylation (Fig. 3d).

**Figure 2.**
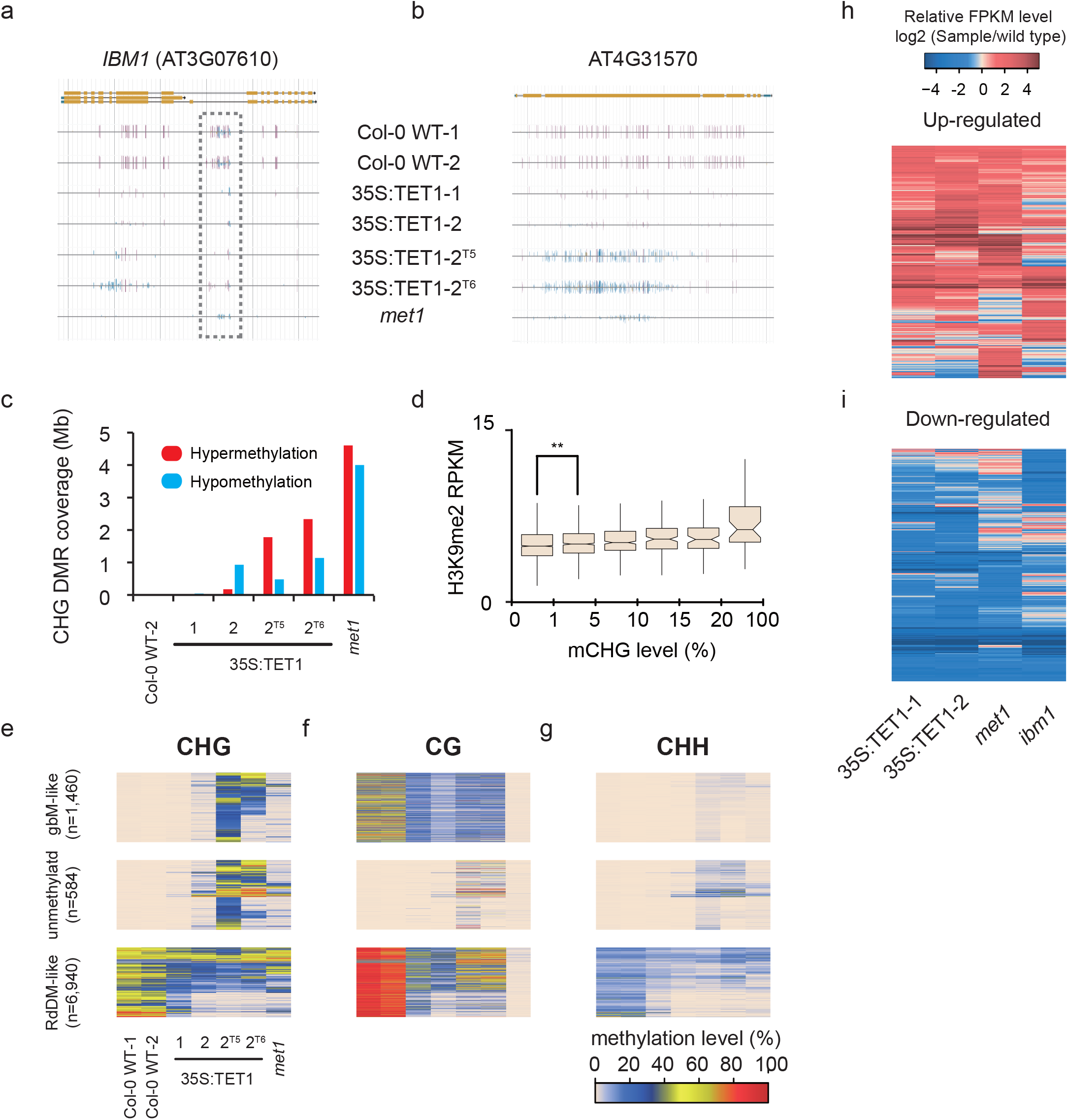
Global fluctuation of CHG methylation in 35S:TET1 plants. (a) Genome browser view of *IBM1* (AT3G07610) in Col-0 WT, three 35S:TET1 transgenic plants and *met1*. A decrease in CG methylation from coding regions was accompanied by an increase in non-CG methylation. Both CG and non-CG methylation was lost from the large intron (purple vertical lines = CG methylation, blue vertical lines = CHG methylation and gold vertical lines = CHH methylation). (b) Genome browser view of a representative CHG hypermethylated region. (c) The amount of the genome affected by differential CHG methylation. These DMRs were defined relative to Col-0 WT-1, as Col-0 WT-2 DMRs were used to assess background interference. (d) Boxplot of H3K9me2 distribution in gbM loci (**, t-test, p-value < 0.01). (e) Heat map of CHG methylation displaying CHG DMRs. Corresponding CG and CHH methylation levels are shown in (f) and (g). Heat maps showing log_2_ transformed FPKM profiles of up-regulated genes (h) and down-regulated genes (i) in two 35S:TET1 transgenic individuals, *met1* and *ibm1* mutants relative to WT.

**Figure 3.**
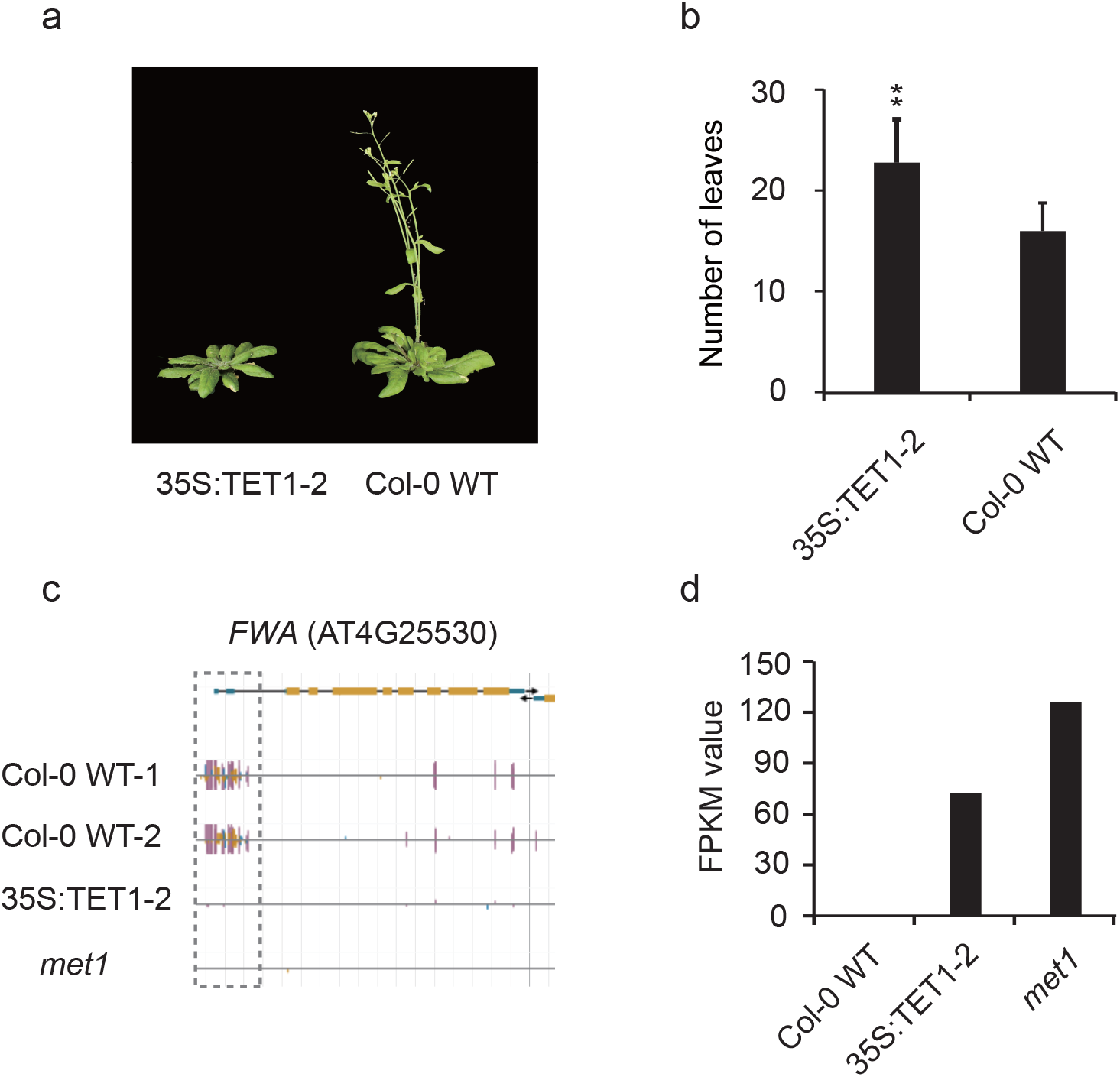
35S:TET1 plants have a delayed flowering phenotype. (a) Photographs of one 35S:TET1-1 transgenic plant and Col-0 WT plant and (b) corresponding number of rosette leaves (**, t-test, p-value < 0.01). (c) Genome browser view of *FWA* (AT4G25530). Both CG and non-CG DNA methylation are depleted from the 5’UTR in 35S:TET1-2 plants (purple vertical lines = CG methylation, blue vertical lines = CHG methylation and gold vertical lines = CHH methylation). (d) Expression level (FPKM) of FWA.

To further characterize regions of differential CHG methylation, identified CHG DMRs in line 35S:TET1-2^T5^ were categorized into discrete groups based on their DNA methylation status in wild-type individuals. Of the 9,917 CHG DMRs identified, 1,460 were in loci that are defined as gbM in wild-type individuals, 584 were in unmethylated regions, and 6,940 of them were in RdDM-like regions (Fig. 2e-g). Interestingly, in line 35S:TET1-2^T5^, 1,408 (96.4%) of the CHG DMRs in gbM-like loci gained CHG hypermethylation, whereas 2,156 (31.1%) of the CHG DMRs in RdDM-like regions lost CHG, in contrast to 595 (8.6%) RdDM-like regions that gained CHG methylation. Lastly, there were 502 (86.0%) loci that are unmethylated in wild-type individuals that gain CHG methylation as well as CG and CHH methylation in the epimutagenized lines (Fig. 2e-g). These results reveal that methods for epimutagenesis can result in both losses and gains in DNA methylation genome-wide.

To characterize the effect of hTET1cd-induced methylome changes on gene expression, we performed RNA-sequencing (RNA-seq) on leaf tissue of wild-type, 35S:TET1-1 and 35S:TET1-2. Compared to wild-type plants, 629 and 736 up-regulated genes were identified in 35S:TET1-1 and 35S:TET1-2, respectively, with 176 and 260 genes overlapping with identified CG DMRs. 1,277 and 1,428 down-regulated genes were identified and 268 and 324 of them overlapped with CG DMRs. There was a high level of overlap in transcriptome changes seen in 35S:TET1-1 and 35S:TET1-2 compared to *met1* and *ibm1* (Fig. 2h **and** i). Of the genes up-regulated in *met1*, 36.8% and 38.7% overlapped with up-regulated genes in 35S:TET1-1 and 35S:TET1-2, respectively (**Supplementary Fig. 2d**). An even greater overlap was observed with down-regulated genes in *met1*, as 60.1% and 65.2% overlapped with down-regulated genes in 35S:TET1-1 and 35S:TET1-2, respectively (**Supplementary Fig. 2e**). These results reveal that hTET1cd expression in *A. thaliana* is a viable approach for accessing hidden sources of allelic variation by inducing expression variation.

### Tet1 expression leads to a delay in the floral transition

In the transgenic plants that were used for WGBS, we observed a delay in the developmental transition from vegetative to reproductive growth (Fig. 3a **and** b). We hypothesized that the observed late-flowering phenotype was associated with the demethylation of the *FWA* (*FLOWERING WAGENINGEN*) locus, as is observed in *met1* mutants^24, 25^. A closer inspection of the DNA methylation status of this locus revealed that DNA methylation was completely abolished, as was methylation at adjacent CHG and CHH sites (Fig. 3c). As in *met1*, the loss of methylation at the *FWA* locus was associated with an increase in *FWA* expression (Fig. 3d), which is known to cause a delay in flowering by restricting the movement of the florigen signal, *FT*, to the shoot apex^26^. These results demonstrate that expression of hTET1cd leads to phenotypic variation by abolishing methylation at some regions in all sequence contexts (CG, CHG and CHH sites).

### Tet1-mediated demethylation can be stably inherited over generations

To assess the stability and inheritance of Tet1-mediated demethylation, T1 individuals expressing 35S:TET1 were self-fertilized, allowing for the loss of the hTET1cd transgene due to allelic segregation. Unexpectedly, transgene-free 35S:TET1-1 T2 individuals (35S:TET1-1.3^−TET1^, 35S:TET1-1.4^−TET1^ and 35S:TET1-1.5^−TET1^) exhibited a reversion to a normal flowering phenotype, and genome-wide methylation levels closely resembled that of wild-type individuals (Fig. 4a). WGBS on T2 individuals retaining the transgene revealed similar levels of CG methylation as the T1 parent (35S:TET1-1.1^+TET1^ and 35S:TET1-1.2^+TET1^). The methylation level of CGs at genes and transposons revealed that demethylation of gbM loci was partially inherited in transgene-free T2 individuals, whereas active remethylation was found at transposons in the same individuals (Fig. 4b-f). These results indicate that an active process likely in the meristem and/or germline is counteracting the activity of the hTET1cd transgene^27^. To quantify how many regions were susceptible to the loss of non-CG methylation, a DMR analysis was conducted for each 35S:TET1-1 T2 individual. 655 and 659 CHG hypomethylated regions were identified in T2 lines retaining the transgene. In contrast, 155, 199 and 201 hypomethylated regions were identified in three transgene-free T2 individuals, respectively (Fig. 4g).

**Figure 4.**
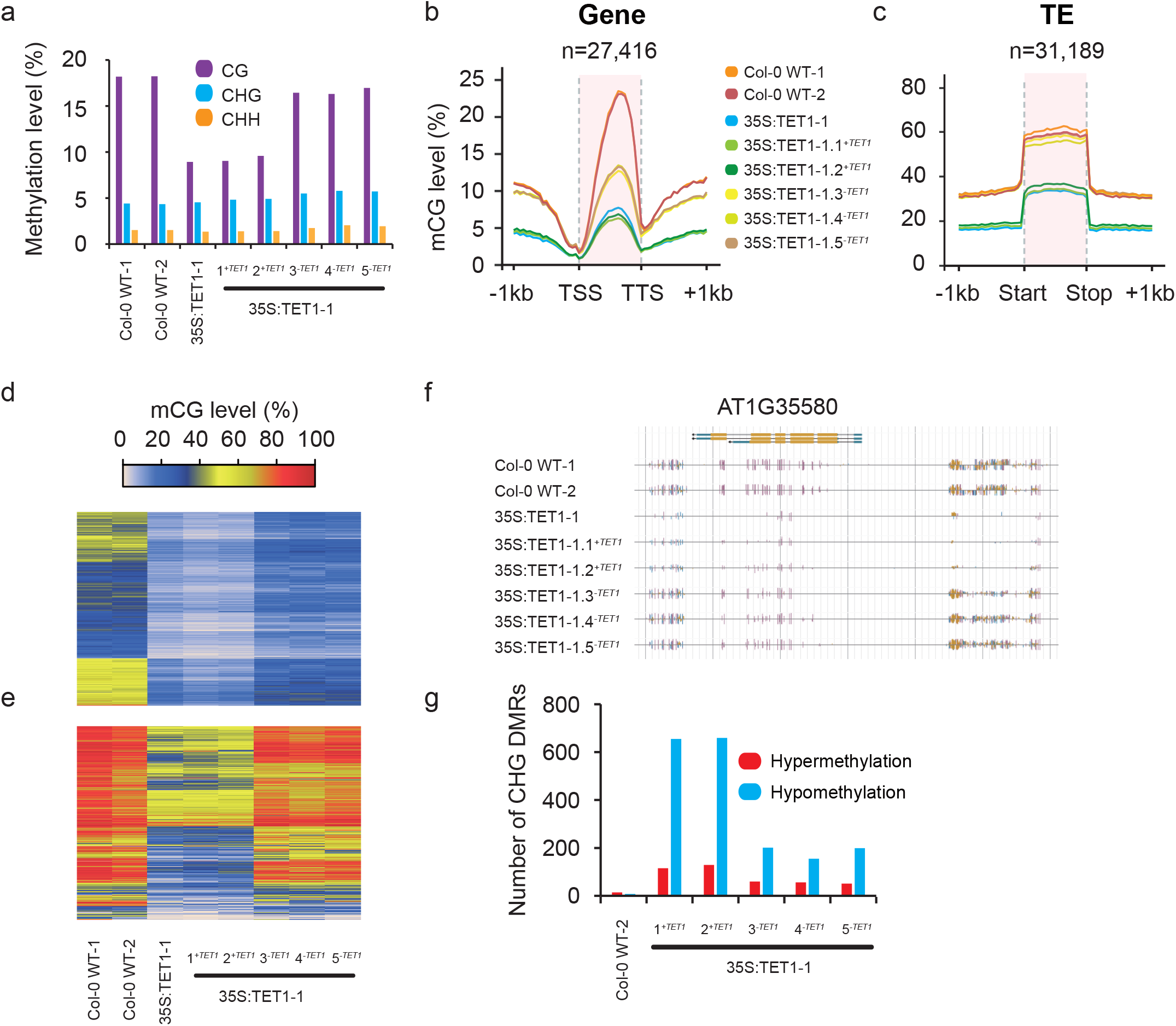
Transgenerational demethylation profile of 35S:TET1 individuals. (a) Bar plots of global methylation levels in two Col-0 WT replicates and 35S:TET1-1 plants. Metagene plots of mCG level across (b) gene bodies and (c) transposable elements. Heat map of mCG level of (d) all gbM genes and (e) transposable elements with >20% mCG in wild type. (f) Genome browser view of a methylome profile of a representative region of the *A. thaliana* genome in 35S:TET1-1 individuals. (g) Number of identified CHG DMRs in ACT2:TET1 T2 plants. These DMRs were defined relative to Col-0 WT-1, as Col-0 WT-2 DMRs were used to assess background.

To determine if increased expression of hTET1cd would increase the likelihood of germline transmittance of demethylation to transgene-free progeny, transgenic *A. thaliana* plants were generated expressing a previously described superfolder GFP (sfGFP) hTET1cd fusion, under control of the *A. thaliana* ACTIN 2 (ACT2) promoter (ACT2:sfGFP-hTET1cd) which is known to have activity in all tissues of juvenile plants, including meristematic tissue^28^. Translation and nuclear localization of the sfGFP-TET1cd fusion protein was confirmed in young cotyledons using confocal microscopy (**Supplementary Fig 3a**). T1 populations transformed with ACT2:sfGFP-hTET1cd exhibited a 27-fold increase in later flowering compared to 35S:hTET1cd, indicating high activity of the sfGFP-TET1cd fusion protein in *A. thaliana* (Fig. 5a). To assess the variation between lines, we performed WGBS on four independent ACT2:TET1 T1 lines (**Supplementary Fig. 4a-c**). Differential levels of global CG demethylation were observed in these four lines, further confirming that plants subjected to epimutagenesis can possess different degrees of demethylation. Subsequent DMR analysis revealed 68,260 CG DMRs, 9,235 CHG DMRs and 2,793 CHH DMRs between these lines, a drastic increase in DMRs compared to those within siblings (Fig. 4g).

**Figure 5.**
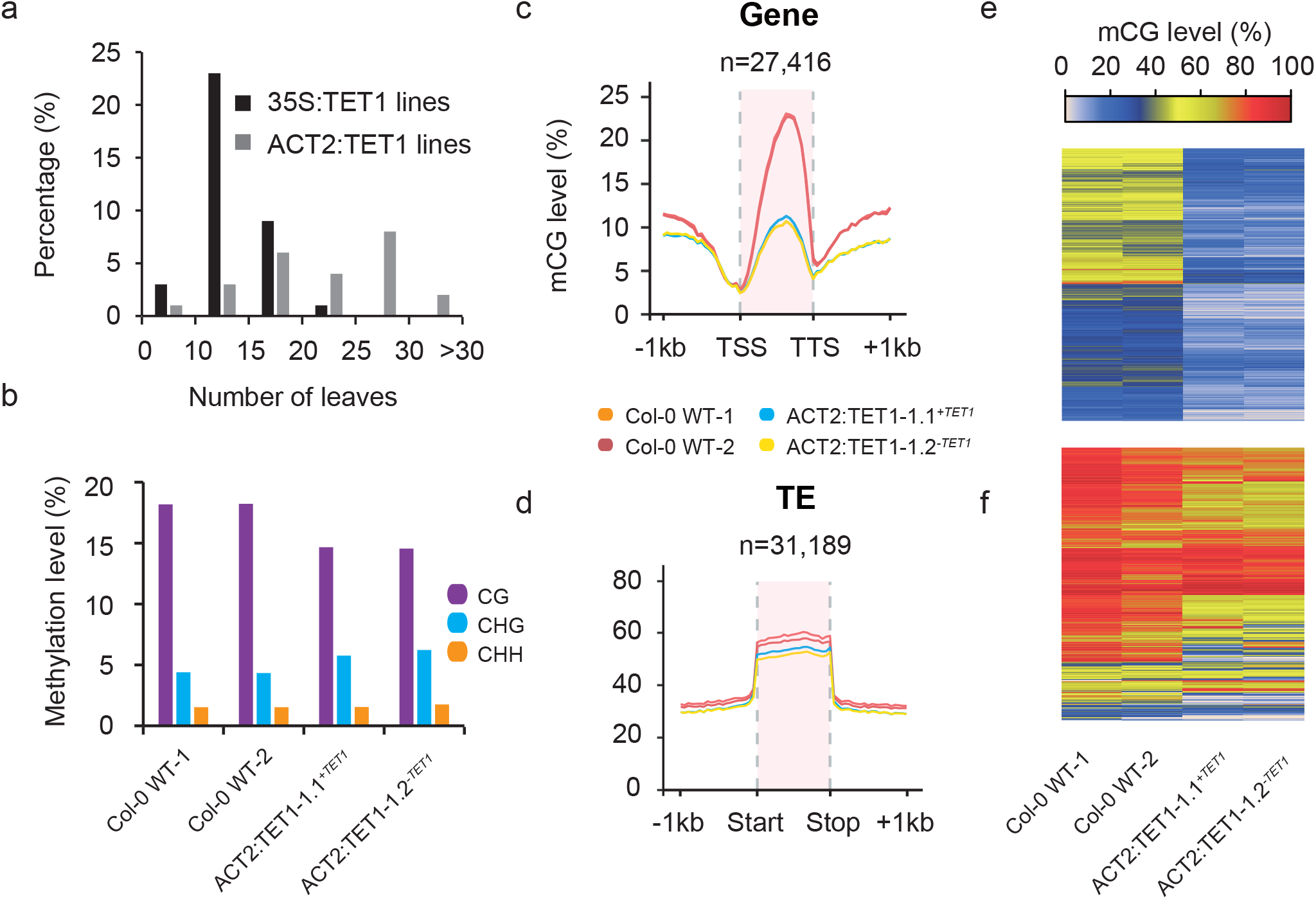
Demethylation profile of ACT2:TET1 individuals. (a) Bar plot of number of rosette leaves in 35S:TET1 and ACT2:TET1 T1 individuals upon flowering. (b) Bar plot of global methylation levels in two Col-0 WT replicates and two ACT2:TET1-1 T2 plants. Metagene plots of mCG level across (c) gene bodies and (d) transposable elements. Heat map of mCG level of (e) all gbM genes and (f) transposable elements with >20% mCG in wild type.

To further assess the inheritance of the demethylation pattern as a result of epimutagenesis, we selected T2 progeny of a late flowering ACT2:TET1-1 T1 individual containing and lacking the transgene. Individuals retaining and lacking the transgene due to allelic segregation both exhibited a late-flowering phenotype, ectopic expression and loss of DNA methylation of *FWA* (**Supplementary Fig 3b and c**). WGBS on these two individuals revealed a reduction in CG methylation that was maintained and stably inherited irrespective of transgene presence, confirming the high activity of demethylation in ACT2:TET1 lines compared to 35S:TET1 lines (Fig. 5b-f). It is unclear exactly why the late-flowering phenotype was stably inherited in the T2 individuals without the transgene in the ACT2 driven lines versus the 35S driven lines, although it is likely a combination of promoter strength and cell type specificity. For stable inheritance of demethylation and the late-flowering phenotype TET1 activity would be required in the meristematic and/or germline cells. Collectively, these results demonstrate the stable inheritance of TET1-mediated demethylation and a delayed floral transition in the absence of the transgene.

## DISCUSSION

The discovery that expression of the catalytic domain of the human TET1 protein in *A. thaliana* leads to widespread loss of CG methylation and enables the creation epimutants without the need for methyltransferase mutants, which often causes lethality in crops. In addition to epimutagenesis, hTET1cd could be used in combination with sequence-specific DNA binding proteins such as dCas9 to direct DNA demethylation in plant genomes, as has been demonstrated in mammalian systems^29, 30, 31, 32, 33^ The stable meiotic inheritance of DNA methylation states in flowering plant genomes provides a stark contrast to the inheritance of DNA methylation in mammalian genomes, where genome-wide erasure of DNA methylation and reprogramming occurs each generation^34^. This property of flowering plant genomes makes them ideal targets of induced-epialleles, as once a new methylation state occurs it is often inherited in subsequent generations. Application of epimutagenesis and the use of TET-mediated engineering of DNA methylation states in economically and agriculturally significant plant species will be an interesting area of future investigation.

## ACKNOWLEDGEMENTS

We thank Nathan Springer for comments and discussions on this study as well as the Georgia Genomics Facility and the Georgia Advanced Computing Resource Center for technical support. This work was supported by the National Science Foundation (MCB-1650331), by The Pew Charitable Trusts and by the Office of the Vice President of Research at UGA to R.J.S. C.H was supported by the National Institutes of Health (NIH HG006827) and is an investigator of the Howard Hughes Medical Institute. W.T.J. was supported by a Scholars of Excellence Fellowship from the University of Georgia and National Institute of General Medical Sciences of the National Institutes of Health award number T32GM007103. The content is solely the responsibility of the authors and does not necessarily represent the official view of the National Institutes of Health.

## AUTHOR CONTRIBUTIONS

R.J.S. conceived the project. L.J., W.T.J. and R.J.S. designed experiments. L.J., W.T.J., X.S., L.H., and C.H. performed research. L.J., W.T.J. and R.J.S. analyzed the data. R.J.S. wrote the paper. All authors discussed the results and commented on the manuscript.

## METHODS

### Synthesis and cloning of the human TET1 catalytic domain

A human TET1 catalytic domain (hTET1cd) sequence (residues 1418-2136) was synthesized by GenScript, and moved to a plant transformation compatible vector (pMDC32) using LR clonase from Life Technologies per the manufacturer’s instructions (Catalog #11791100). ACT2:sfGFP-hTET1cd was subcloned by Genscript in the pMDC32 vector background using the sfGFP-TET1cd fragment from Addgene plasmid #82561. The ACT2 promoter sequence was kindly provided by Dr. Richard Meagher.

### Plant transformation and screening

The hTET1cd sequence in the pMDC32 vector was transformed into *Agrobacterium tumefaciens* strain C58C1 and plated on LB-agar supplemented with kanamycin (50 μg/mL), gentamicin (25 μg/mL), and rifampicin (50 μg/mL). A single kanamycin resistant colony was selected and used to start a 250-mL culture in LB Broth Miller liquid media supplemented with gentamicin (25 μg/mL), kanamycin (50 μg/mL), and rifampicin (50 μg/mL), which was incubated for two days at 30°C. Bacterial cells were pelleted by centrifugation at 4,000 RPM for 30 minutes and the supernatant decanted. The remaining bacterial pellet was re-suspended in 200 mL of 5% sucrose with 0.05% Silwet L77. Plant transformation was performed using the floral dip method described by Clough and Bent^35^. Seeds were harvested upon senescence and transgenic plants were identified via selection on ½ LS plates supplemented with Hygromycin B (25 μg/mL). 35S:TET1-1 is a T1 individual, 35S:TET1-2 is a T3 plant, 35S:TET1-3 is a T4 plant. All transgenic individuals chosen for analysis contain independent insertions of hTET1 cd and are not the result of single-seed decent unless otherwise noted.

### DNA and RNA isolation

*A. thaliana* leaf tissue was flash-frozen and finely ground to a powder using a mortar and pestle. DNA extraction was carried out on all samples using the DNeasy Plant Mini Kit (QIAGEN), and the DNA was sheered to approximately 200 bp by sonication. RNA was isolated from finely ground flash-frozen leaf tissue using Trizol (Thermo Scientific). For RT-qPCR, RNA was further treated with TURBO™ DNase (Thermo Scientific) according to manufacturers instructions. One μg of RNA was subsequently reverse transcribed with M-MuLV Reverse Transcriptase according to manufacturers instructions (NEB). RT-qPCR was used to analyzed cDNA populations using PP2AA-3 (AT1G13320) as an endogenous control, and was performed on a Roche LIghtCycler 480 instrument using SYBR Green detection chemistry. The genes assayed by this method were *IBM1-S*, *IBM1-L*, and FWA. Primers used for RT-qPCR were designed using PrimerQuest from Integrated DNA Technologies (www.idtdna.com/PrimerQuest/). Primers sequences used for RT-qPCR: PP2A-F: 5’ AATGAGGCAGAAGTTCGGATAG 3’, PP2A-R: 5’CAGGGAAGAATGTGCTGGATAG 3’, ibm1s-F: 5’ TCTTTCTTCTAAGTCTGTCCATTCT 3’, ibm1s-R: 5’ GTGACCGATTAGGAAATGGTATCT 3’, ibm1L-F: 5’ CCGAAGCCAAAGTGGAGATA 3’, ibm1L-R: 5’ CTTCCTCTTCCGTAGACTTCTTT 3’, FWA-F: 5’ CAAGAT GGTGGAAGGATGAGAA 3’, FWA-R: 5’ CTCTGTTCTTCAGTGGGATGAG 3’.

### Library construction

Genomic DNA libraries were prepared following the MethylC-seq protocol without use of the bisulfite conversion step. MethylC-seq libraries were prepared as previously described in^36^. RNA-seq libraries were constructed using Illumina TruSeq Stranded RNA LT Kit (Illumina, San Diego, CA) following the manufacturer’s instructions with limited modifications. The starting quantity of total RNA was adjusted to 1.3 μg, and all volumes were reduced to a third of the described quantity. TAB-seq libraries were prepared as previously described in^16^.

### ChIP library preparation

Leaves were treated with formaldehyde to covalently link protein to DNA, washed several times with distilled water, patted dry and ground into fine powder in liquid nitrogen. Chromatin was extracted with a series of extraction buffers and sonicated. The final chromatin solution was incubated overnight with anti-H3K9me2 antibody (Cell Signaling Technology, 9753S) coated Dynabeads protein A (Life Technologies, 10002D) to precipitate the immune complex. After a few washes, the immune complex was eluted and incubated at 65 degrees in the presence of a high concentration of NaCl in a water-bath overnight. After degrading the proteins with proteinase K, DNA was recovered by phenol/chloroform/isoamyl alcohol extraction followed by ethanol precipitation. The DNA pellet was then dissolved in 30μl of nuclease-free water.

### Sequencing

Illumina sequencing was performed at the University of Georgia Genomics Facility using an Illumina NextSeq 500 instrument. For MethylC-seq and TAB-seq, raw reads were trimmed for adapters and preprocessed to remove low quality reads using cutadapt 1.9.dev1^37^. For RNA-seq and ChIP-seq, these processes were carried out by Trimmomatic v0.32^38^. The mutant allele of *met1* is *met1-3* and methylome data was downloaded under accession GSE39901. The mutant allele of *ibm1* is *ibm1-6*.

### MethylC-seq data processing

Qualified reads were aligned to the *A. thaliana* TAIR10 reference genome as described in^39^. Chloroplast DNA (which is fully unmethylated) was used as a control to calculate the sodium bisulfite reaction non-conversion rate of unmodified cytosines. All conversion rates were >99% (**Supplementary Table 1**). The list of gbM genes used in this study was previously curated^18^. Heat maps were clustered by Complete Linkage method conducted by R (https://www.r-project.org). All methylation levels reported in all analyses are presented as differences in absolute values, including defining DMRs and calculating hyper/hypomethylated regions. The only exception is in the comparison of mCG loss between gbM, where we used a percentage difference.

### RNA-seq data processing

Qualified reads were aligned to the *A. thaliana* TAIR10 reference genome using TopHat v2.0.13^40^ (Supplementary Table 2). Gene expression values were computed using Cufflinks v2.2.1^41^. Genes determined to have at least 2-fold log_2_ expression changes by Cufflinks and passed tests were identified as differentially expressed genes.

TAB-seq data processing. Qualified reads were aligned to the *A. thaliana* TAIR10 reference genome using Methylpy as described in^39^. A modified lambda DNA sequence was used to assess the quality of prepared libraries. In the added lambda sequence, all non-CG cytosines are unmethylated and CG cytosines are methylated to 5mC. The ‘non-conversion’ rate is used to measure the rate of non-CG cytosines failing to be converted to thymines after bisulfite treatment. The ‘5mC non-conversion’ is used to estimate the 5mCG cytosines failing to be converted to thymines after TET treatment. (**Supplementary Table 3**).

### ChIP-Seq Data Analysis

Qualified reads were aligned to the *A. thaliana* TAIR10 reference genome using Bowtie 1.1.1 with following parameters: bowtie -m 1 -v 2 --best --strata --chunkmbs 1024 -S. Aligned reads were sorted using samtools v 1.2 and clonal duplicates were removed using SAMtools version 0.1.19 (**Supplementary Table 4**).

### Metaplot analysis

For metaplot analyses, twenty 50 bp bins were created for both upstream and downstream regions of gene bodies/TEs. Gene bodies/TE regions were evenly divided into 20 bins. Weighted methylation levels were computed for each bin as described previously^42^.

### DMR analysis

Identification of DMRs was performed as described in^43^ and adjusted p-value (Benjamini-Hochberg correction) 0.05 was adopted as the cutoff. Only DMRs with at least 5 DMSs (Differential Methylated Sites) and a 10% absolute methylation level difference within each DMR were reported and used for subsequent analysis. For coverage calculations, each sample was combined with two Col-0 WT replicates to identify DMRs. Each sample was compared with both Col-0 WT replicates separately and for a DMR to be identified it must have been identified in both comparisons. Absolute methylation differences of +/− (50% for CG, 10% for CHG and CHH) were defined as hyper/hypo methylation, respectively. DMRs overlapping regions with mCG >= 5%, mCHG and mCHH >= 1% in both two Col-0 WT replicates were defined as RdDM-like regions. DMRs overlapping regions with mCG >= 5%, mCHG and mCHH < 1% in both two Col-0 WT replicates were defined as gbM regions. DMRs overlapping regions with all three contexts less methylated at less than 1% in both Col-0 WT replicates were defined as unmethylated regions. Overlap comparisons were performed using bedtools v2.26.0^44^.

### Data availability

The data generated from this study has been uploaded to Gene Expression Omnibus (GEO) database and can be retrieved through accession number GSE93024.

